# Antidepressant-like effects of P2 purinergic antagonist PPADS is dependent on serotonergic and noradrenergic integrity

**DOI:** 10.1101/086983

**Authors:** Cassiano R. A. F. Diniz, Murilo Rodrigues, Plínio C. Casarotto, Vítor S. Pereira, Carlos C. Crestani, Sâmia R. L. Joca

## Abstract

Depression is a common mental disorder affecting around 350 million of individuals globally. The available antidepressant drug monotherapy is far from ideal since it has an efficiency of approximately 60% and takes around 3-4 week to achieve clinical improvement. Attention has been paid to the purinergic signaling regarding neuropathological mechanisms, since it might be involved in psychiatric disorders, such as depression. In fact, blockade of purinergic P2X receptors induces antidepressant-like effects in preclinical models. However, the mechanisms involved in this effect are not yet completely understood. The present work investigated the interplay between a P2X receptor antagonist (PPADS) and clinically used antidepressant drugs on the forced swimming test, an animal model predictive of antidepressant effect. We observed significant synergistic effect of PPADS combined with sub-effective doses of fluoxetine or reboxetine in the FST. Moreover, depletion of serotonergic or noradrenergic systems, with PCPA and DSP-4 treatment, respectively, blocked the antidepressant-like effect of PPADS. No increase in locomotion, a possible source of confusion on FST data, was detected in any of the treated groups. Our results indicate the antidepressant-like effect of PPADS depends on the integrity of serotonergic and noradrenergic transmission.

## 1. Introduction

The World Health Organization (WHO) estimates that around 350 million of individuals are affected by depression globally, and it predicts it will be the main cause of morbidity and loss of productivity among all health conditions in 2030 (World Health Organization, 2016). The current antidepressant drug therapy, based on increasing monoamines availability (Elhwuegi, 2004), is only effective in approximately 60% of the patients, and takes 3 to 4 weeks to be clinically effective (Blier, 2003; Racagni and Popoli, 2008).

Great attention has been paid to the purinergic signalling in neuropathological disorders, since several studies have shown purinergic involvement in multiple neurodegenerative, neurological and psychiatric diseases such as Alzheimer, Parkinson, anxiety, schizophrenia, drug addiction, and, of importance to the present work, depression, (Abbracchio et al., 2009; Burnstock, 2008). Purinergic neurotransmission emerged from the work of prof. Burnstock describing adenosine 5´-triphosphate (ATP) as non-adrenergic, non-cholinergic inhibitory transmitter in the guinea-pigs (Abbracchio and Burnstock, 1994; Burnstock, 1972; Burnstock, 1976). Since then, ATP has been described as a cotransmitter in noradrenergic (Sperlagh et al., 1998), gabaergic (Jo and Role, 2002), glutamatergic (Mori et al., 2001) and cholinergic (Richardson and Brown, 1987) synaptic terminals, but vesicles containing exclusively ATP have also been reported (Pankratov et al., 2006). ATP released in synaptic cleft can be degraded by ectonucleotidases into various metabolites, such as adenosine, which also acts as a ligand for purinergic receptors (Zimmermann, 2006). Purinergic signaling is also implicated in signal integration between glial cells and neuronal synapses (tripartite synapse), inasmuch as ATP is recognized as a neurotransmitter accessible to perisynaptic glia and neurons, in addition to being released from glial and neuronal cells (Pascual et al., 2005; Araque et al., 2014; Halassa and Haydon, 2010; Lalo et al., 2014; Kato et al., 2004; Koizumi et al., 2005).

Purinergic neurotransmission comprises receptors for adenosine and ATP, respectively termed P1 and P2. Based on molecular cloning and pharmacological differences, P2 receptors can be divided into P2Y and P2X receptors (Abbracchio and Burnstock, 1994). P2Y are G-protein coupled receptors that modulate the inositol -1,4,5-triphosphate (IP3) and the dyacil-glycerol (DAG) levels from membrane phosphoinositide metabolism (Pfeilschifter, 1990; Burnstock, 2007), while P2X are ion channels permeable to Na^+^, K^+^ and Ca^2+^ (Burnstock, 2007; Benham and Tsien, 1987; Bean, 1992). Currently, 8 subtypes of metabotropic P2Y receptors (P2Y_1,2,4,6,11,12,13,14_) and 7 subtypes of ionotropic P2X receptors (P2X_1-7_) have been described (Burnstock, 2007). P2 receptors are widely expressed in the central nervous system (CNS), either in glial or neuronal cells (Burnstock, 2008), including cerebral structures involved in emotional behavior such as hippocampus, cerebral cortex, ventral tegmental area and locus coeruleus (Norenberg and Illes, 2000).

Recently, our group reported that the P2X receptor antagonist isoPPADS showed antidepressant-like effect associated to a reduction in nitric oxide production in the prefrontal cortex, in animals submitted to the forced swimming test (Pereira et al., 2013). More recently, another group reported that chronic treatment with a P2X7 antagonist induced antidepressant-like effect in the chronic mild stress model of depression (Iwata et al., 2016). However, there is no evidence addressing if the antidepressant-like effect induced by P2 receptor antagonists involves serotonergic and/or noradrenergic mechanisms. Monoamine and purinergic interplay is plausible, given that the second messenger NO coming from glutamate/ATP interaction might control the production and stability of serotonin (Kuhn and Arthur, 1997; Fossier et al., 1999), and that P2X7R knockout animals showed changed cerebral levels of noradrenaline and serotonin (Csolle et al., 2013).

Therefore, the present work aimed at investigating the interplay of a P2X receptor antagonist and clinically effective antidepressant drugs on the forced swimming test, an animal model predictive of antidepressant effect. In order to address to do that combined treatment of P2 antagonist (PPADS) with either selective serotonin reuptake inhibitor or selective noradrenaline reuptake inhibitor was used to discern ATP/serotonin or ATP/noradrenaline interplay. Furthermore, the effect of DSP-4, a selective neurotoxin to noradrenergic terminals; or PCPA, a tryptophan hydroxylase inhibitor, on PPADS effects was addressed.

## 2. Material and methods

### 2.1 Animals

A total of 352 male Swiss mice weighing 30-40g were used. The animals were housed in groups of 6 to 8 per cage (570 cm^2^) in a temperature controlled room (24±1 ºC) under standard laboratory conditions with free access to food and water and a 12 h light/12 h dark cycle (light on at 6:30 a.m.). Animals were randomly assigned for the experimental groups described bellow and procedures were performed in conformity with the Brazilian Council for the Control of Animals under Experiment (CONCEA), which comply with international laws and politics. The local Ethical Committee approved the experimental protocol (146/2009), and all efforts to minimize animal suffering were made.

### 2.2 Drugs

Pyridoxalphosphate-6-azophenyl-2′,4′-disulfonic acid tetrasodium salt (PPADS, a non-selective P2R antagonist, Tocris, USA, #0625), fluoxetine hydrochloride (FLX, selective serotonin reuptake inhibitor, Prati-Donaduzzi, Brazil, #713198), reboxetine mesylate (RBX, selective noradrenaline reuptake inhibitor, Ascent Scientific, USA, #157), p-Chlorophenylalanine (PCPA, tryptophan hydroxylase inhibitor, Sigma-Aldrich, EUA, #C6506), N-(2-chloroethyl)-N-ethyl-2-bromobenzylamine hydrochloride (DSP4, noradrenergic neurotoxin, TOCRIS, USA, #2958) were used. All drugs were prepared administered intraperitoneally (i.p.) at 10 mL/Kg, except for DSP4, which was infused intracerobroventricularly in 1µl volume. DSP4 and FLX were dissolved in Tween 80 2% in sterile saline, while all other drugs were dissolved in sterile isotonic saline. 2,2,2-tribromoethanol (Aldrich Chemical, USA, #T48402) was used for stereotaxic surgery and chloral hydrate (Sigma Aldrich, USA, #V000554) was used to euthanize the animals for tissue collection.

### 2.3 Apparatus and procedure

#### Forced Swimming Test (FST)

Animals were individually submitted to 6 min of forced swimming in glass cylinders (height 25 cm, diameter 17 cm) containing 10 cm of water at 24 – 26°C. The test was videotaped and immobility time (characterized by limb movements necessary for floating) was measured during the last 4 minutes of the test by a trained observer that was blinded to the treatment condition. All behavioral procedures are in accordance to established protocols (Porsolt et al., 1977). The water was changed after each trial (Abel and Bilitzke, 1990).

#### Open Field Test (OFT)

The OFT was used to measure locomotor activity of the animals according to Zanelati et al (Zanelati et al., 2010). Mice were placed individually in a circular open field arena (40 cm in diameter and 50 cm high Plexiglas wall) for 6 min. The exploratory activity was videotaped and the number of crossings between the quadrants of the arena was measured during the last 4 minutes, by an observer that was blinded to the treatment condition. After each trial, the arena was cleaned with 70% alcohol solution.

#### Sample collection and high-performance liquid chromatography (HPLC)

Animals were deeply anesthetized with chloral hydrate 5% (0.75g/kg, ip) and the brains removed. The frontal cortex and hippocampus were dissected on ice and processed for determination of serotonin, noradrenaline and serotonin metabolite (5HIAA) levels as described by Patel et al. (2005). The samples were homogenized in 0.1 M perchloric acid, afterward centrifuged at 13000rpm for 10 min at 4 °C. The supernatant was automatically injected into the chromatographic system to quantify neurotransmitters and metabolites by electrochemical detection in high-performance liquid chromatography (HPLC). The chromatograph (Waters® Alliance) consisted of a stripping column Symmetry® C18, 5μm (150x4.6 mm) and mobile phase flow 1ml/min with the following composition: buffer (88.2%; citric acid 0.05 M, sodium octyl sulfate 250mg/L, EDTA 0.1mM, potassium chloride 2mM and pH adjusted to 3.2 with NaOH), methanol (9.3%) and acetonitrile (2.5%). The mobile phase was vacuum filtered and degassed ultrasonically. The calibration curve was constructed with standard solutions of 2.5, 5, 10, 25, 50, 100 and 200ng/ml of norepinephrine, serotonin and 5HIAA, which were injected into the chromatograph in triplicate. The detection and limit of quantification were 0.64 and 2.13ng/ml for serotonin; 1.72 and 5.73ng/ml for 5HIAA and 0.87 and 2.88ng/ml for noradrenaline. Samples with concentrations found below the limit of quantitation were discarded. Finally, the concentrations of the substances were corrected by the mass of the dissected tissue samples being expressed in ng of substance per mg of tissue.

#### Stereotaxic surgery and intracerebral administration

Mice were anesthetized with 2,2,2-tribromoethanol 2.5% (10ml/Kg intraperitoneal, ip) and fixed in a stereotaxic frame. One Stainless steel guide cannula (0.7 mm OD) was implanted and aimed at the lateral ventricle (coordinates: anteroposterior= −0.3mm from lambda, lateral= 1.2mm, dorsoventral= 2.5mm) according to the Paxinos and Franklin’s atlas (Paxinos and Franklin, 1997). The cannula tip was placed 1mm above the site of injection and attached to the skull bone with stainless steel screws and acrylic cement. A stylet inside the guide cannula prevented obstruction.

Five to seven days after the surgery, intra-cerebroventricular (i.c.v), the injections were performed with a thin dental needle (0.3 mm OD). A volume of 1 μl per animal was injected in 30s using a micro-syringe (Hamilton) controlled by an infusion pump (Insight Equipamentos Científicos, Brazil). A polyethylene catheter (PE10) was interposed between the upper end of the dental needle and the micro-syringe. The movement of an air bubble inside the polyethylene catheter confirmed drug flow.

#### Histology

After the behavioral tests, the mice were deeply anesthetized with chloral hydrate 5% (0.75g/kg, ip) and a dental needle was inserted through the guide cannulae to infuse 0.5µl of methylene blue. Immediately after, the animals were euthanized and the brains were removed. The injection sites were confirmed as correct, on fresh brain, after the dye has spread to the whole brain through cerebroventricular way. Results from injections outside the target area were discarded from statistical analysis.

### 2.4 Experimental design

#### Experiment 1: Effective and sub-effective doses of FLX, RBX and PPADS on FST

Experimentally naïve mice received a single ip injection of PPADS (3, 6.25, 12.5mg/kg), FLX (10, 20, 30 mg/kg), RBX (2.5, 5, 10 mg/kg) or vehicle and, 30min later, submitted to FST. An independent cohort of animals received single ip injections of effective doses of PPADS (6.25mg/kg), FLX (20mg/kg), RBX (5mg/kg) or vehicle and submitted to OFT 30min later.

#### Experiment 2: Add on effect of PPADS sub-effective dose to FLX or RBX sub-effective doses on FST

Independent group of animals received two injections. The first injection was saline, FLX 10mg/kg or RBX 2.5mg/kg, immediately followed by a second injection of saline or PPADS 3mg/kg. Thus, six groups were obtained: VEH+VEH, FLX+VEH, RBX+VEH, PPADS+VEH, FLX+PPADS, RBX+PPADS. The animals were submitted to FST 30min after the last administration. An independent cohort of animals received ip injections of FLX (10mg/kg), RBX (2.5mg/kg) or vehicle, immediately followed by PPADS (3mg/kg), and submitted to OFT 30min after the last administration.

#### Experiment 3: Effect of 5-HT depletion over PPADS effects on FST and OFT

Experimentally naïve mice received daily ip injections of PCPA (150mg/kg for 4 days) or VEH. On the fourth day, PCPA or VEH were administered 30min before VEH or PPADS (6.25mg/kg) administration. The animals were submitted to FST 30min after the last injection. An independent cohort of animals was submitted to the same treatment regimen described above and submitted to the OFT 30min after the last injection.

#### Experiment 4: Effects of NA depletion over PPADS effects on FST and OFT

Experimentally naïve mice, with surgically implanted cannula, received i.c.v. infusion of VEH or DSP4 (20µg) followed, 24h later, by ip injections of PPADS (6.25mg/kg) or VEH, and submitted to FST 30min after ip administrations. An independent cohort of animals was submitted to the same treatment regimen described above and submitted to the OFT 30min after the last injection.

#### Experiment 5: 5HT, NA and 5HIAA levels on PFC and HPC of PCPA and DSP4 treated animals

Experimentally naïve mice received daily ip injections of PCPA (150mg/kg for 4 days) or VEH and euthanized 30min after the last administration. An independent cohort of mice, received icv infusion of DSP4 (20ug) or VEH 24h before euthanasia. The levels of 5HT, NA and 5HIAA levels were analyzed on PFC and HPC through HPCL as described.

### 2.5 Statistical analysis

Data from experiment 1 and experiment 2 (OFT) were analyzed by one-way ANOVA, followed by Dunnett’s *post hoc* test when appropriate. Experiments 2 (FST), 3 and 4 were analyzed by two-way ANOVA, with injections as factors, followed by Fisher’s LSD *post hoc* test when appropriate. Data from experiment 7 was analyzed by Student’s t test. Statistical differences were considered significant when p<0.05.

## 3. Results

### Experiment 1: Effective and sub-effective doses of FLX, RBX and PPADS on FST

One-way ANOVA indicates a significant effect of drug treatment on immobility time in the FST [F_9,60_=3.453. p<0.05]. Treatment with PPADS (6.25mg/kg), FLX (20mg/kg) and RBX (5mg/kg) significantly decreased immobility time on FST (Dunn’s *post hoc* test), as seen in Figure 1A. Effective doses of PPADS (6.25mg/kg), FLX (20mg/kg) and RBX (5mg/kg) were not able to change locomotor behavior of animals tested on OFT (Figure 1B - F_3,20_=1.715, non-significant – NS).

**Fig 1.**
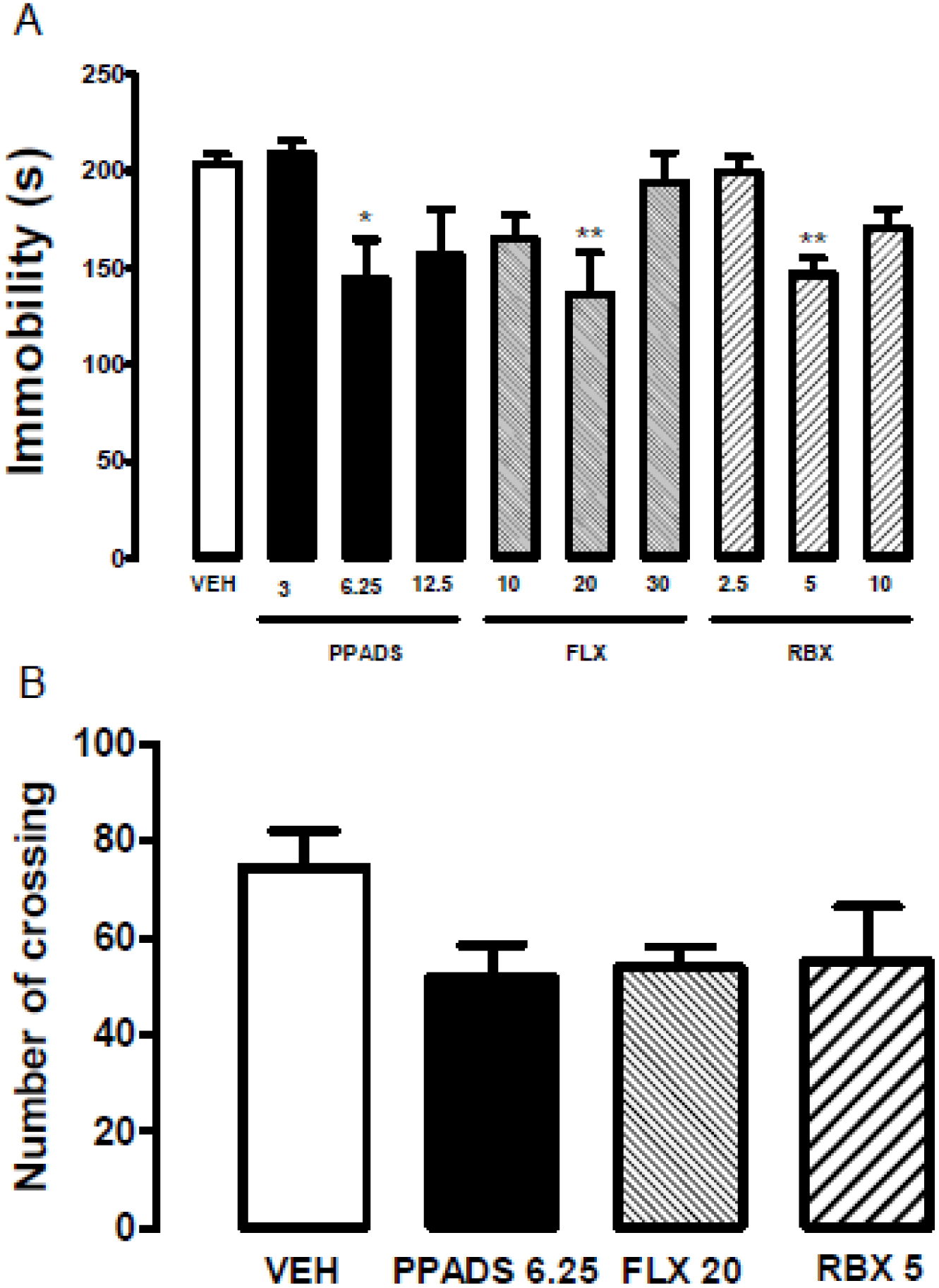
Effect of three different doses (mg/kg) of PPADS, FLX or RBX on FST and OFT. (A) PPADS (3, 6.25, 12.5), FLX (10, 20, 30), RBX (2.5, 5, 10) or vehicle (10mL/kg) were administered 30min before FST (n=6,7,7,7,7,5,5,7,10,9, respectively). Data are expressed as mean ± SEM of immobility time (s); *p<0.05, **p<0.01 from control group. (B) PPADS (6.25mg/kg, n=6), FLX (20mg/kg, n=6), RBX(5mg/kg, n=6) or VEH (n=6) was administered 30min before being submitted to OFT. Data are expressed as Mean ±SEM of total quadrant traveled in the OFT.

### Experiment 2: Add on effect of PPADS sub-effective dose to FLX or RBX sub-effective doses on FST

Two-way ANOVA indicates a significant effect of first injection [F_2,63_=5.697; p<0.01] and second injection [F_1,63_=6.255; p<0.05], but no interaction between them [F_2,63_=1.281; NS]. Combination of sub-effective doses of PPADS (3mg/kg) and FLX (10mg/kg) or RBX (2.5mg/kg) significantly decreased immobility time during FST (Figure 2A - F_5,69_=4.243. p<0.05). Combination of sub-effective doses of PPADS (3mg/kg) and FLX (10mg/kg) or RBX (2.5mg/kg) was also not able to change locomotor activity of animals on the OFT (Figure 2B - F_2,14_=0.6730, NS).

**Fig 2.**
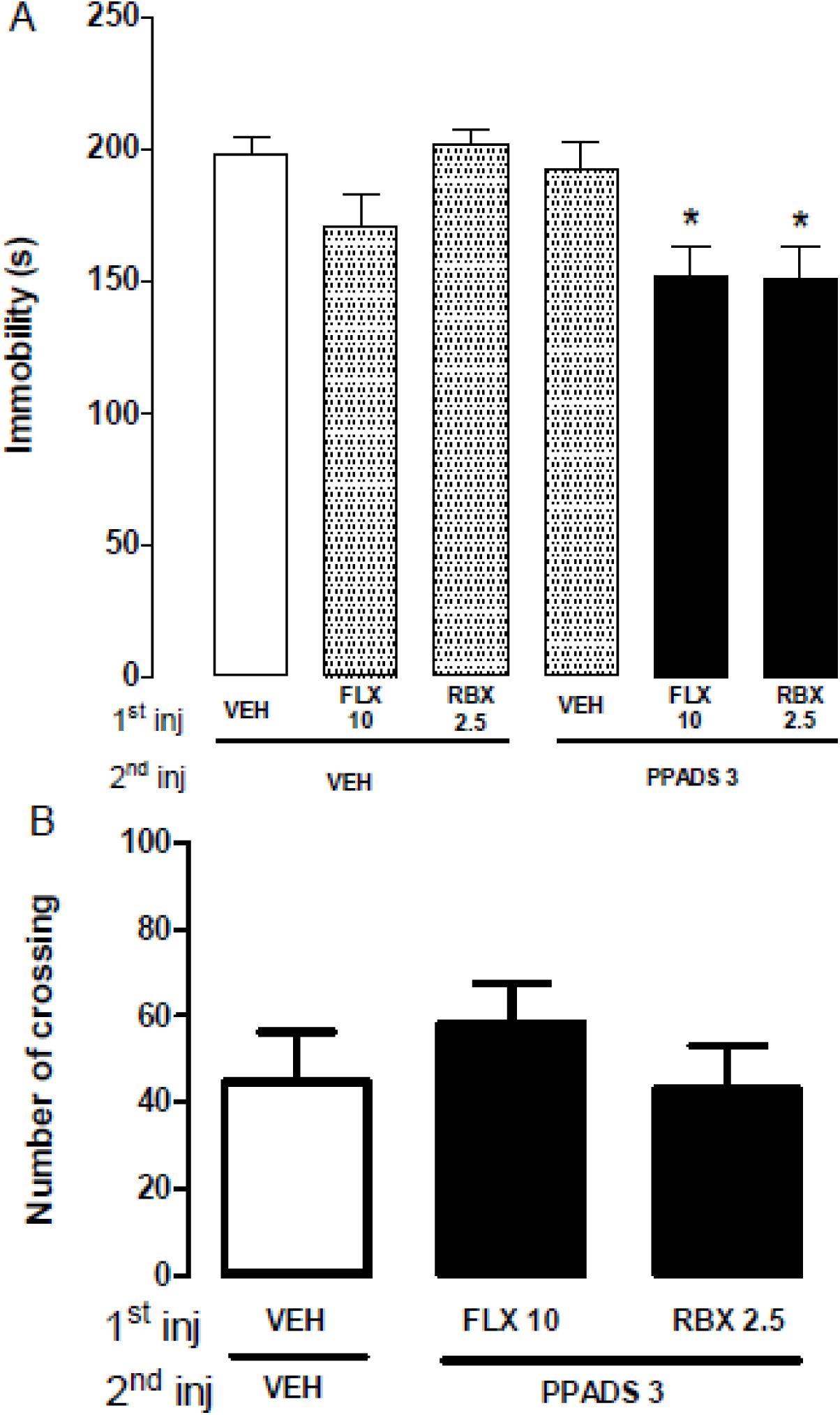
Add on effect of PPADS sub-effective dose with FLX or RBX sub-effective doses on FST and OFT. (A) Sub-effective dose of FLX (10mg/kg) or RBX (2.5mg/kg) was carried out after PPADS sub-effective dose (3mg/kg) administration, 30min before FST. Six groups were obtained: VEH + VEH (n=16), FLX + VEH (n=14), RBX + VEH (n=6), VEH + PPADS (n=16), FLX + PPADS (n=15), RBX + PPADS (n=8). Data are expressed as mean ± SEM of immobility time (s); *p<0.05 from control group. (B) Sub-effective dose of FLX (10mg/kg) or RBX (2.5mg/kg) was carried after PPADS sub-effective dose (3mg/kg) administration, 30min before OFT. Three groups were obtained: VEH + VEH (n=6), FLX + PPADS (n=6), RBX + PPADS (n=5). Data are expressed as Mean ±SEM of total quadrant traveled in the OFT.

### Experiment 3: Effect of 5-HT depletion over PPADS effects on FST and OFT

Two-way ANOVA indicated a significant effect of second injection [F_1,27_=8.64; p<0.01] but not first injection [F_1,27_=0.25; NS] or interaction between factors [F_1,27_=2.115; NS]. Pretreatment with PCPA was able to block PPADS effect on FST [F_3,27_=3.631, p<0.05 - Figure 3A]. No locomotor change was induced by any of the treatments (first injection: F_1,19_=0.859; NS; second injection: F_1,19_=1.316; NS; interaction: F_1,19_=0.466; NS - figure 3B).

**Fig 3.**
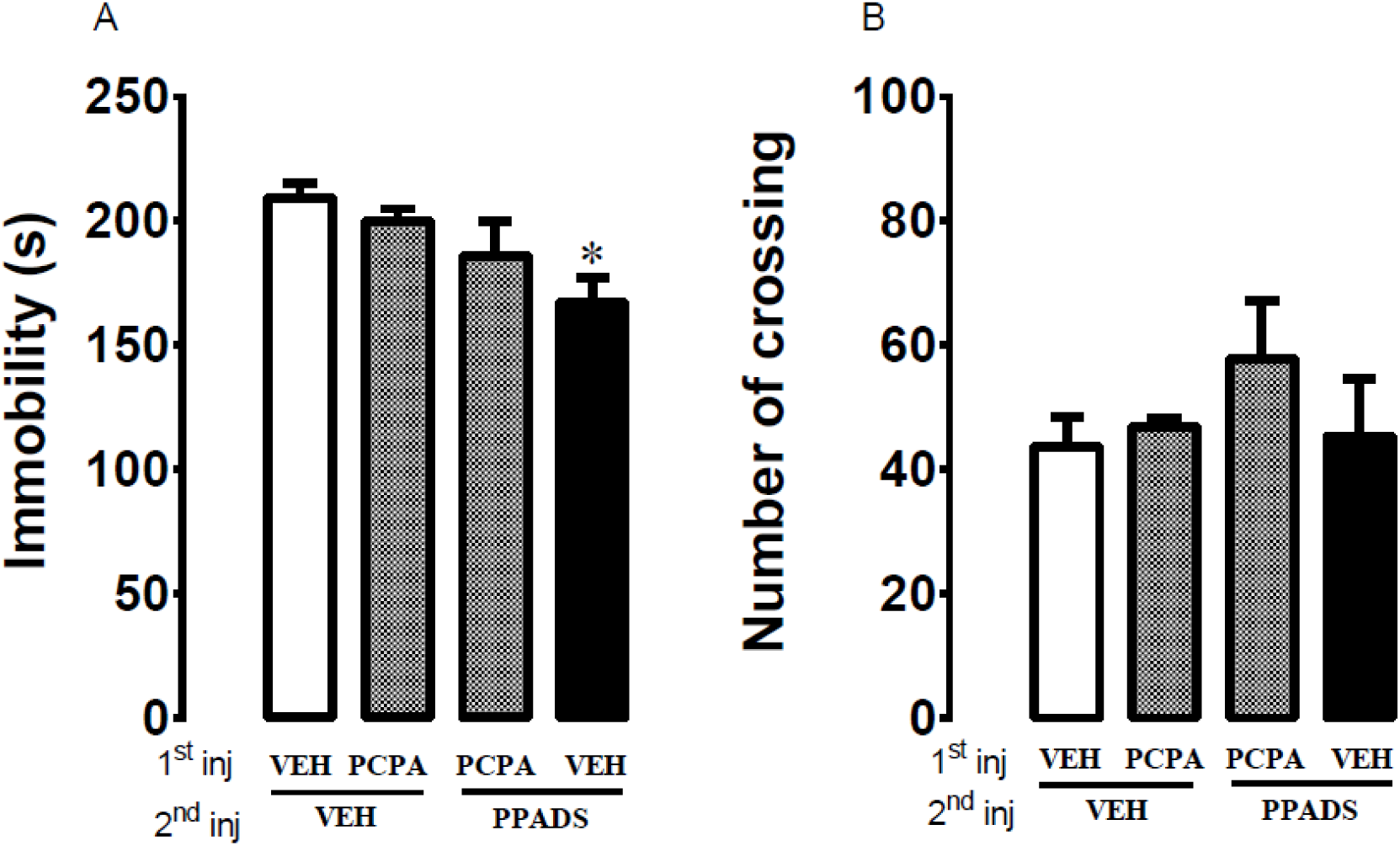
Effect of 5-HT depletion over PPADS effects on FST and OFT. (A) PCPA (150mg/kg) was administered once a day for 4 days. On the fourth day, PCPA or VEH were administered 30min before VEH or PPADS (6.25mg/kg). The animals were submitted to the FST 30min after the last injection. Four groups were obtained: VEH + VEH (n=7), PCPA + VEH (n=8), PCPA + PPADS (n=8), VEH + PPADS (n=8). Data are expressed as mean ± SEM of immobility time (s). (B) The same treatment protocol was established as described above, but animals were submitted to the OFT instead of the FST. Four groups were obtained: VEH + VEH (n=6), PCPA + VEH (n=6), PCPA + PPADS (n=5), VEH + PPADS (n=6). Data are expressed as Mean ±SEM of total quadrant traveled in the OFT. *p<0.05 from control group.

### Experiment 4: Effect of NA depletion over PPADS effects on FST and OFT

Two-way ANOVA indicates a significant interaction between first and second injections [F_1,31_=8.624; p<0.01], however no effect of them isolated [first injection: F_1,31_=1.016; second injection: F_1,31_=1.198, NS for both]. Pretreatment with DSP4 was able to prevent PPADS effect on FST [F_3,31_=3.423, p<0.05 – Figure 4A]. In addition, a significant effect of the DSP4 infusion on OFT was observed [Figure 4B - F_1,24_=9.580; p<0.01].

**Fig 4.**
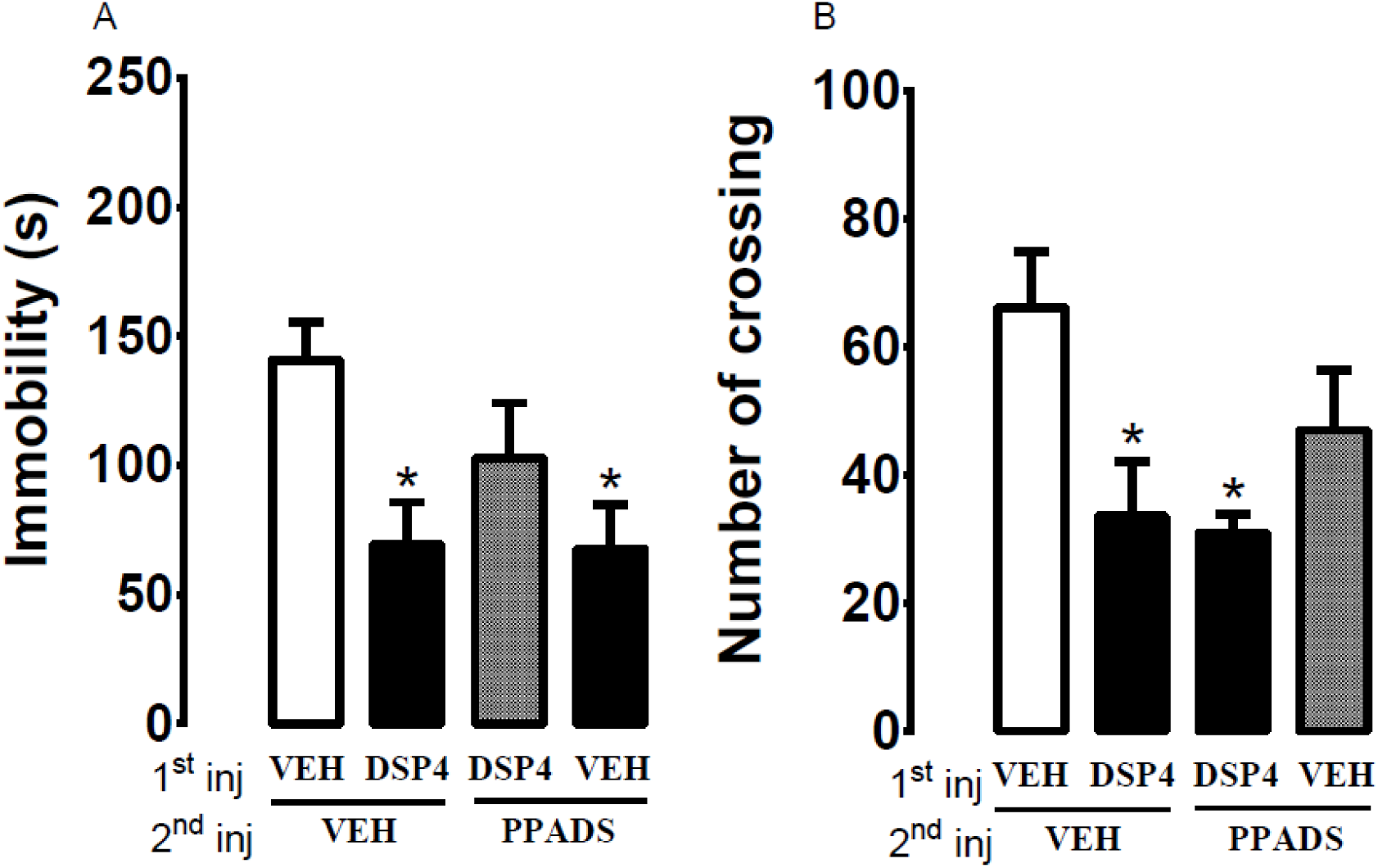
Effect of NA depletion over PPADS effects on FST and OFT. (A) DSP4 (20µg) was infused i.c.v. 24h before FST. Injection of PPADS (6.25mg/kg) was performed 30min before FST. Four groups were obtained: VEH + VEH (n=7), DSP4 + VEH (n=8), DSP4 + PPADS (n=9), VEH + PPADS (n=8). Data are expressed as mean ± SEM of immobility time (s). (B) The same treatment protocol was established as described above, but animals were submitted to the OFT instead of the FST. Four groups were obtained: VEH + VEH (n=7), DSP4 + VEH (n=7), DSP4 + PPADS (n=7), VEH + PPADS (n=7). Data are expressed as Mean ±SEM of total quadrant traveled in the OFT. *p<0.05 from control group.

### Experiment 5: 5HT, NA and 5HIAA levels on PFC and HPC of PCPA and DSP4 treated animals

According to student’s t test there was a significant decrease of NA (t_8_=2.451), 5HIAA (t_8_=2.491) and 5HT (t_8_=3.885) hippocampal levels in DSP4 treated mice. No difference was observed with DSP4 treatment in frontal cortex (NA t_10_=0.018; 5HIAA t_10_=1.178 and 5HT t_10_=1.672), see figure 5A. Student´s t test identified a reduction in NA (t_7_=2.488) frontal cortex levels from PCPA treatment but no change was observed in 5HIAA (t_6_=1.100) and 5HT (t_7_=1.096) levels. In hippocampus, 5HIAA (t_7_=2.647) levels were significantly reduced by PCPA treatment, but no change was observed in NA (t_7_=0.159) and 5HT (t_7_=0.490) levels, as found in figure 5B.

**Fig 5.**
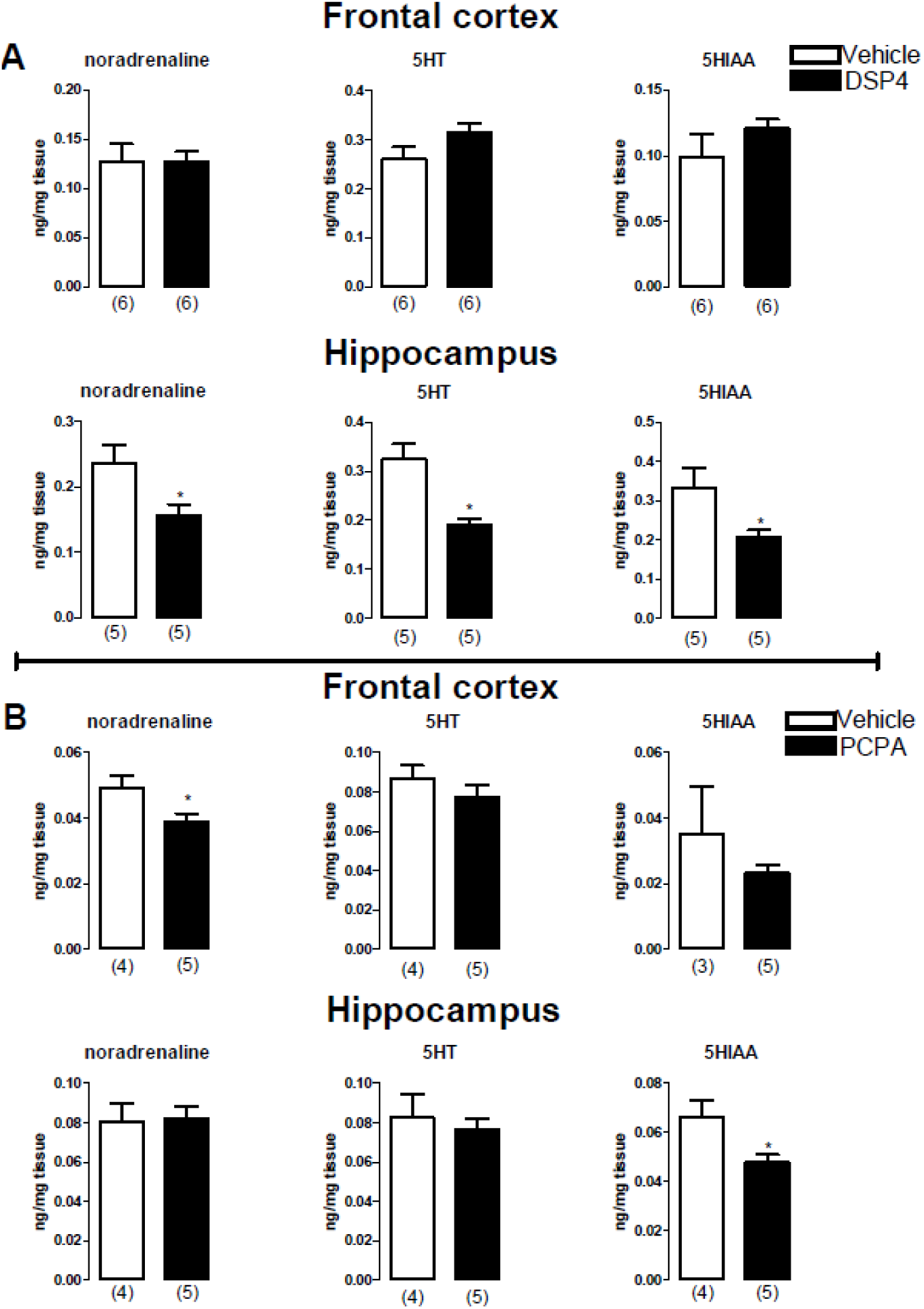
5HT, NA and 5HIAA levels on PFC and HPC of PCPA and DSP4 treated animals. (A) DSP4 was infused i.c.v. 24h before euthanasia. Two groups were obtained: VEH (n=6) and DSP4 (n=6). (B) PCPA (150mg/kg) was administered once a day for 4 days. On the fourth day PCPA was administered 1 hour before euthanasia. Serotonin, noradrenaline and serotonin metabolite levels were analyzed on frontal cortex and hippocampus with HPCL method. All data are expressed as Mean ±SEM of ng/mg tissue, *p<0.05 from control group. Number of animals per group here is based in number of samples which HPCL MS determination was possible (in the graphic), although number of animals euthanized is higher (VEH n=6/DSP4 n=6; VEH n=5/PCPA n=5).

## 4. Discussion

This work replicated our previous finding that PPADS treatment induces dose-dependent antidepressant-like effect in mice submitted to FST. Based on that, it was possible to evaluate a putative add on effect of combining sub-effective doses of PPADS with sub-effective doses of FLX or RBX in animals submitted to FST. Results evidenced a synergistic effect of PPADS co-treatment with FLX and RBX, thus suggesting the involvement of monoaminergic mechanisms. Accordingly, PCPA and DSP4 pretreatment, which decreased serotonin and noradrenaline levels, prevented the antidepressant-like effect of PPADS. In addition, none of these effects can be the result of unspecific locomotor effects, since none of the changes is likely as confounding factor.

Stressful events, a key factor for susceptibility to depression, would be expected to affect behavior in the FST. In fact, immobility time of rats exposed to swim session is increased after uncontrollable stress and decreased by antidepressant treatment (Porsolt et al., 1977; Cryan et al., 2002; Nestler and Hyman, 2010). Although there is no consensus about what immobility in FST represents, a behavioral despair or a passive-coping (Cryan et al., 2002), the fact that it is decreased by all known antidepressants confers to it a good predictive validity and allows the study of mechanism of action of current and putative new drugs. In this work, systemic treatment with P2R antagonist reduced immobility time in the FST, in a dose-dependent fashion, as earlier observed (Pereira et al., 2013), further supporting the involvement of the purinergic system as a new target for putative antidepressant drugs. This has been recently corroborated by another study where chronic blockade of P2X7 receptors induced antidepressant-like effects in the chronic mild stress model (Iwata et al., 2016).

ATP signaling may work as a neurotransmitter and even as a gliotransmitter (Pascual et al., 2005; Araque et al., 2014; Halassa and Haydon, 2010). This dual functional role of the purinergic system contributes to the communication between astrocytes and synaptic terminals, as a tripartite synapse (Pascual et al., 2005). ATP could evoke astrocyte calcium waves leading to the release of other transmitters such as glutamate, GABA and glycine (Guthrie et al., 1999). ATP could also be exclusively released from central synapses (Pankratov et al., 2006) or co-released from glutamatergic (Mori et al., 2001), noradrenergic (Sperlagh et al., 1998), gabaergic (Jo and Role, 2002) and cholinergic (Richardson and Brown, 1987) synaptic terminals. The interaction between glutamatergic and purinergic neurotransmission has been the most studied so far, with ATP facilitating neuronal and astrocyte glutamate release (Burnstock, 2002; Burnstock, 2011; Shen et al., 2005). Glutamate, as well as NO, which arises from NMDAR activation, can control serotonin production and stability (Kuhn and Arthur, 1997; Fossier et al., 1999). Evidence indicates that ATP itself is also able to attenuate serotonin release in rat brain cortex via P2R activation (Von Kügelgen et al., 1997). Conversely, serotonin is also able to modulate ATP levels in the brain (Koren-Schwartzer et al., 1994). In this context, the add-on effect observed with concomitant PPADS and FLX administration could derive from P2R blockade favoring serotonergic availability. In a previous work from our group, PPADS was able to attenuate NO levels in the prefrontal cortex at the same dose that induced antidepressant-like effects in the FST (Pereira et al., 2013). Decreasing NO synthesis in the brain produces antidepressant-like effects (Diniz et al., 2016), which are dependent on serotonin levels (Harkin et al., 2003). Altogether, this evidence suggests serotonin involvement in PPADS-induced antidepressant-like effects. In line with this possibility, PCPA pretreatment was able to block the effect of PPADS on FST.

HPLC analysis revealed that PCPA treatment decreased noradrenaline levels in the frontal cortex, while it decreased serotonin main metabolite, 5-hydroxyindoleacetic acid (5HIAA), levels in hippocampus. Changes in brain catecholamines after PCPA systemic administration, although subtle, were observed in the original work of Koe and Weissman (1966). Tyrosine hydroxylase was also slightly inhibited in vitro with PCPA infusion and plasma tyrosine levels were slightly reduced after systemic PCPA injection (Koe and Weissman, 1966). Curiously, 5HT levels did not change in either frontal cortex or hippocampus, which is in disagreement with other works (Shutoh et al., 2000; Fletcher et al., 2001; Page et al., 1999). Differences in methodological analysis of serotonin content or in drug administration and brain dissection protocols might explain such controversy. Besides that, other structures involved with the modulation of the behavioral consequences of stress such as the hypothalamus, bed nucleus of the stria terminalis, nucleus accumbens and amygdala could present serotonin levels changes, but this possibility was not verified in the present work. Although no change was observed on serotonin levels after PCPA treatment, 5HIAA hippocampal levels were decreased. Concentration of 5HIAA has been used to estimate the activity of serotonergic neurons (Shannon et al., 1986). In fact, electrical stimulation of the dorsal raphe nucleus increased 5HIAA levels and the 5HIAA/5-HT concentration ratio in the nucleus accumbens, amygdala, suprachiasmatic nucleus and dorsomedial nucleus (Shannon et al., 1986). Therefore, PCPA treatment used in the present work, although slightly, was able to alter 5HT and noradrenaline neurochemistry in the brain, what might have contributed to its blocking effect on PPADS behavioral effects in mice FST. Similar PCPA treatment protocols have been used by others and described its effectiveness in blocking antidepressant effects in the FST (Harkin et al., 2003).

Our results also suggest a synergistic antidepressant-like effect with concomitant RBX and PPADS. Activation of presynaptic P2R is shown to increase the release of noradrenaline in the locus coeruleus (Tschopl et al., 1992), whereas it decreases noradrenaline availability in the rat cortex (Von Kügelgen et al., 1994) and hippocampus (Koch et al., 1997), thus showing an important and complex role for ATP in regulating noradrenaline levels in the brain. Our results showing that combination of sub-effective doses PPADS and RBX induced synergistic effects suggests the involvement of central noradrenergic mechanisms. The fact that DSP4 was able to suppress PPADS effects gives further support to that possibility.

As shown in our results, DSP4 treatment decreased not only noradrenaline levels in the hippocampus, but also serotonin (5HT) and 5HIAA levels. In the frontal cortex, no change was observed on noradrenaline, serotonin or 5HIAA. Since the hippocampus is located close to the ventricular wall, it is possible that i.c.v. drug infusion reached hippocampal region easier than more distant areas, such as the frontal cortex. The main DSP4 neurotoxic mechanism of action involves noradrenergic neurotransmission, although one work has reported decreased serotonin levels in the cerebellum of rat brains and in the spinal cord (Jonsson et al., 1981), which is in agreement with our results. Several reports describe noradrenaline levels decrease in cerebral regions after systemic injections of DSP4, whereas many other reports from microdialysis state that extracellular noradrenaline levels are actually increased (for review see Ross and Stenfors, 2015). Therefore, the antidepressant-like effect of DSP4 *per se* could be explained by increased extracellular levels of noradrenaline in some particular brain regions, although the overall levels of noradrenaline are decreased (as shown in HPLC results). In this sense, the absence of effect from a combined use of PPADS and DSP4 could be the result of a complex interplay between both drugs acting through the whole CNS. In a speculative way, the combined action of the drugs is perhaps dependent of noradrenaline levels available to be released and/or dependent of serotonin levels in the hippocampus or from other areas not studied.

Regarding DSP4 experiments, a decreased locomotor activity was verified in both DSP4 + VEH and DSP4 + PPADS groups, which might originate from DSP4 effect *per se* on locomotor activity. In fact, it cannot be excluded that the reversal of PPADS antidepressant-like effect by DSP4 might be related to a non-specific DSP4 effect on locomotion. However, that is unlikely, provided that DSP4’s effect in decreasing locomotor activity did not prevent its own antidepressant-like effect.

In conclusion, the present study further corroborate that the blockade of P2R induces antidepressant effects and suggests the involvement of serotonin and noradrenaline release in structures related to stress adaptation and depression, such as the hippocampus and the cerebral frontal-cortex. Nevertheless, the functional role of purinergic transmission in depression neurobiology is still poorly understood and more studies regarding the purinergic/monoaminergic interplay are necessary to better understand the psychopathological mechanisms involved on that.

## 5. Acknowledgements

The authors would like to thank to Flavia F. Salata and Elisabete Zocal Paro Lepera for her technical support. This work was supported by research grants from FAPESP and CNPq. C.R.F. Diniz received a PhD fellowship from FAPESP.

### 6. Conflict of interest

The authors declare no conflict of interest.

